# Protein-DNA target search relies on Quantum Walk

**DOI:** 10.1101/2020.10.14.337840

**Authors:** M. D’Acunto

## Abstract

Protein-DNA interactions play a fundamental role in all life systems. A critical issue of such interactions is given by the strategy of protein search for specific targets on DNA. The mechanisms by which the protein are able to find relatively small cognate sequences, typically 15-20 base pairs (bps) for repressors, and 4-6 bps for restriction enzymes among the millions of bp of non-specific chromosomal DNA have hardly engaged researcher for decades. Recent experimental studies have generated new insights on the basic processes of protein-DNA interactions evidencing the underlying complex dynamic phenomena involved, which combine three-dimensional and one-dimensional motion along the DNA chain. It has been demonstrated that protein molecules spend most of search time on the DNA chain with an extraordinary ability to find the target very quickly, in some cases, with two orders of magnitude faster than the diffusion limit. This unique property of protein-DNA search mechanism is known as *facilitated diffusion*. Several theoretical mechanisms have been suggested to describe the origin of facilitated diffusion. However, none of such models currently has the ability to fully describe the protein search strategy.

In this paper, we suggest that the ability of proteins to identify consensus sequence on DNA is based on the entanglement of π-π electrons between DNA nucleotides and protein amino acids. The π-π entanglement is based on Quantum Walk (QW), through Coin-position entanglement (CPE). First, the protein identifies a dimer belonging to the consensus sequence, and localize a π on such dimer, hence, the other π electron scans the DNA chain until the sequence is identified. By focusing on the example of recognition of consensus sequences by EcoRV or EcoRI, we will describe the quantum features of QW on protein-DNA complexes during search strategy, such as walker quadratic spreading on a coherent superposition of different vertices and environment-supported long-time survival probability of the walker. We will employ both discrete- or continuous-time versions of QW. Biased and unbiased classical Random Walk (CRW) has been used for a long time to describe Protein-DNA search strategy. QW, the quantum version of CRW, have been widely studied for its applications in quantum information applications. In our biological application, the walker (the protein) resides at a vertex in a graph (the DNA structural topology). Differently to CRW, where the walker moves randomly, the quantum walker can hop along the edges in the graph to reach other vertices entering coherently a superposition across different vertices spreading quadratically faster than CRW analogous evidencing the typical speed up features of the QW. When applied to protein-DNA target search problem, QW gives the possibility to achieve the experimental diffusional motion of proteins over diffusion classical limits experienced along DNA chains exploiting quantum features such as CPE and long-time survival probability supported by environment. In turn, we come to the conclusion that, under quantum picture, the protein search strategy does not distinguish between one-dimensional (1D) and three-dimensional (3D) case.

**Significance:** Most biological processes are associated to specific protein molecules binding to specific target sequences of DNA. Experiments have revealed a paradoxical phenomenon that can be synthesized as follows: proteins generally diffuse on DNA very slowly, but they can find targets very fast overwhelming two orders of magnitude faster than the diffusion limit. This paradox is known as *facilitated diffusion*. In this paper, we demonstrate that the paradox is solved by invoking the quantum walk picture for protein search strategy. This because the protein exploits quantum properties, such as long-time survival probability due to coherence shield induced by environment and coin-position entanglement to identify consensus sequence, in searching strategy. To our knowledge, this is the first application of quantum walk to the problem of protein-DNA target search strategy.

## 1. Introduction

The ability of DNA-binding proteins to locate specific binding sites within the genome constitutes a fundamental basis of many biological processes based on target search strategy [1–3]. This ability to bind selectively to DNA enables cell processes linked to RNA transcription, DNA packing, replication, recombination or repair. The understanding of molecular recognition process underlying the specific protein-DNA binding selection is hence extremely important in structural biology. Protein features involved in binding selection to DNA, including direct (base) or indirect (shape) readout, can be classified as: i) *Enzymatic*, where the main function of the protein is to modify DNA. ii) *Transcription Factor* (TF), where the main function of the protein is to regulate transcription and gene expression and, in turn, iii) *Structural-DNA Binding Proteins*, where the main function of the protein is to support DNA structure, DNA bending or to aggregate other proteins [4–5]. Recently, the interest in the target search has greatly increased because of the possibility of single-molecule experiments [6–8]. Such experiments have evidenced that during the search the protein molecule experiences an alternating searching dynamics alternating freely 3D diffusing behaviour around the DNA chain and scanning 1D motion along DNA chains [9–12]. Since the seminal work of Riggs *et al*. [13], who firstly evidenced that *lac* repressor is capable to bind to its target sites *in vitro* with rates overwhelming the 3D diffusion limit (~10^8^-10^9^ M^−1^s^−1^), many experimental studies involving conditions *in vitro*, *in silico* and *in vivo* have suggested the idea commonly recognized of *facilitated diffusion* for protein-DNA searching process [6–8, 14–17]. In the simplest form, the facilitated diffusion consists of cycling DNA filaments by proteins through non-specifically bound states, where the protein is bound to non-specific DNA and free states where the protein is not associated to DNA. However, in the picture of facilitated diffusion, since the previous description could provide searching processes inherently slow, to locate and bound to the target site, the proteins scanning a DNA chain must interrogate multipole sites in a single association event through two diffusion-based mechanisms that are the sliding and hopping mechanics [3, 18–21]. In addition, a third mechanism by which proteins may move from one site to another is the so-called intersegmental transfer, which can be considered a sort of combination of sliding-hopping mechanisms. For example, TFs not only to be able to locate their targets, wherewith have a high equilibrium binding probability, in short time, generally less than few minutes. This search strategy requires that in their nonspecific binding mode, TFs although strongly associated with the DNA must be able to diffuse randomly along the genome. Standard 1D diffusion is not able to describe the short time involved in TFs search strategy on bacterial genome of length *N*≈5×10^6^ bps (≈150-170μm). With a 1D typical diffusion constant of *D*_1_≈1μm^2^/s, one would get a time *t*~*N*^2^/*D*_1_~10^4^s for a single TF. Analogously, maximal rate obtained by Debye-Smoluchowski with a protein diffusion coefficient of *D*_3_~10μm^2^/s provides a value *k*_*DS*_=4π*D*_3_*a*_*0*_~10^8^M^−1^s^−1^, two orders of magnitude lower than the observed value, *a*_*0*_=0.34nm is the protein-DNA distance at the binding site [22].The facilitated diffusion model is currently the most accepted theory of DNA target searching [20, 22–26] and also the model able to explain the greatest number of experimental results. Such model is an interplay of 3D and 1D diffusion processes. Firstly, a protein diffuses in a 3D space filled by DNA filaments with which the protein is found to experience a non-specific interaction. The protein can switch to 1D diffusion along a DNA filament, where the protein can encounter the specific target or dissociates from the DNA filament due to its finite dissociation constant. The model assumes that the faster-than-3D diffusion binding is achieved by exploiting a *reduction of dimensionality* of the cognate site search process reducing the search to a 1D diffusion. The interplay of relative times spent in 3D or 1D diffusion motion when the sliding distance is short, ~50 bps, is the strength of the model. The complex protein dynamics, other than 3D diffusion, involves hopping, jumping, intersegmental transfer behaviours featured in one of the three states [19]: (i) an unbound state performing diffusion motion including jumping to DNA chain, (ii) a search state, weakly bounded to the DNA chain, performing one dimensional sliding motion, (iii) a recognition state, tightly bounded to the DNA. Although facilitated diffusion explains the different time scale involved in the recognition process, however, currently, there is a relevant discrepancy between experiments and theoretical models on protein-DNA site identification and conformational transitions between searching conformations. Many experimental studies have demonstrated that many proteins associate to specific sites on DNA in times shorter than predicted from three-dimensional bulk solution calculations [27–29]. Another critical limitation of facilitated diffusion is that some specific mechanisms of protein-DNA binding and recognition mechanisms are totally disregarded. For example, the energetic nature of the contact between proteins and DNA, that today is still not clarified. Contacts between DNA and proteins are generally noncovalent, traditionally classified as hydrogen bonding (water-mediated), ionic (salt bridges), van der Waals, and hydrophobic interactions. It has been evidenced that protein-DNA complexes can vary the leading role of the different forces for the contact [5]. For 129 protein-DNA complexes, van der Waals interactions seem play a major role with respect to hydrogen bonds. On the contrary, other 139 protein-DNA complexes have evidenced a major role of hydrogen bonding with respect to van der Waals, hydrophobic interactions or electrostatic interactions [5]. However, there is a common consensus on the fundamental role of nucleobase-amino acid π-π contacts [30]. The large number of π-π interactions between the DNA nucleobases and Tyr, Phe, His or Trp in 141 DNA-protein complexes, and today it is commonly recognized that the proximity of nucleobases and aromatic amino acids implies that aromatic-aromatic interactions, both π-π or C/N-H…π, play a leading role in stabilizing protein-DNA complexes and supporting nucleic acid recognition both on base or shape readout.

In this paper, we suggest that π-π interaction in protein-DNA complexes shows quantum entanglement during the nucleobases recognition through a position base entanglement alongside the sequence identification as typically evidenced in QW. We apply for the first time QW to the problem of protein-DNA search problem, where the DNA is assumed to be a linear, compact polymer generated by four monomers, A, C, G and T. It is well known that theoretical and experimental studies have evidenced that protein-DNA searching processes exploit CRW to achieve facilitated diffusion [17, 25]. Our effort will be to show that if quantum version of CRW is used to describe the complex processes during protein-DNA interaction, the diffusional experimental results can be immediately obtained when the protein (the walker) exploits the speed-up properties of QW and peculiar entanglement-based behaviour enforced by the environmental fluctuations. Recently, there is a growing interest in the application of quantum properties, long-lived quantum coherence, superposition, tunneling and entanglement to biological systems [31–33], ranging from enzyme catalysis [34–35], to photosynthesis (although this is a controversial subject) [36–38], to the implication of quantum entanglement in avian navigation [39], to quantum tunneling in olfaction [40], genetic mutation [41], while speculative suggestions pay attention on the link between quantum coherence and consciousness [42].

QW is the quantum equivalent to CRW [43]. On the contrary, to CRW, where usually the walker explores the space randomly, in quantum case, the walker resides at a vertex on a graph and is allowed to move or to hop along the edge in the graph to get to other vertices. One fundamental difference, between quantum and classical picture, is that in the quantum case the walker coherently enters a superposition across different vertices at any step. In our QW, the protein is the walker and the DNA chain the graph, and the nucleotides the vertices. Even when CRW and QW take place on the same structure, classical and quantum walks can display drastically different behaviour. For example, while classical walk loses memory of the starting site, QW exhibits transition probabilities, which depend on the starting site. In addition, classical mean-square displacement of the walk up to time *t* changes as <*x*^2^(*t*)>~*t*^*β*^, where the diffusion exponent β can assume the *standard* value (β=1), or the anomalous value (β≠1). Quantum transport relies generally on a speed-up ballistic power-law as <*x*^2^(*t*)>~*t*^*α*^, where generally α≅2, but can reach α~3 in non-Markovian approaches [44]. Even when classical anomalous diffusion takes a value ~2 for the β exponent, the classical and quantum behaviours of the walker are strongly different. In the presence of a chaotic structure quantum wave propagation can exhibit localization, a quantum property that can be exploited in a target search and delocalization of wave packet spreads quadratically. In Appendix A it is demonstrated the quadratic speed-up ability of continuous time QW on a tight-binding Hamiltonian describing the search of targets on long DNA chains by the proteins walker.

In discrete QW, the Hamiltonian requires a Coin operator. The changes on the spreading behaviour of the walker under general conditions show the entanglement between the coin state and the walker position. This entanglement is a consequence of the ability of Coin operator at each step to control the interference pattern of the walker. This entanglement, known as CPE, depends strongly on the initial position. By using CPE, the protein can detecting a sequence by identifying an initial dimer of the sequence and subsequently explore the nucleotides around the dimers. In turn, another basic advantage for a fast sequence identification of DNA sequences by exploiting QW is the positive interference of environment contrasting decoherence. The environment is represented, for example, by the stabilizing role of aqueous solution containing Na^+^ and Mg^+^ contrasting the negative charge of phosphate groups and vibration modes of DNA operate as decoherence shield [45].

We suggest that exploration of DNA sequences by proteins is based on the fundamental role of charge transfer due to π-electrons where the protein works as a quantum walker exploiting a typical feature of QW, such as CPE, between initial states identified by the protein-DNA contact, where initial states are given by specific dimer nucleotide bases of the sequence to be identified. Quantum properties are needed to achieve to typical experimental rates, because diffusion rate of π-electrons injected by the protein as donor states at room temperatures is approximately *D*_1_ ≅ *kλ*^2^ ≅ 10^2^ μm^2^/s, in agreement with experimental results on EcoRV [27], where *k*≅10^7^Hz is donor-acceptor charge transfer at physiological conditions and 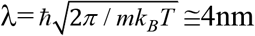 is the de Broglie length of π-electrons at room temperatures, *k*_*B*_ is the Boltzmann constant and *T* the temperature. Analogously, proteins bind to DNA with a time *t*_0_=10^−8^s and with a diffusion *D*=10μm^2^/s, a protein can scan a length *l*=4(*Dt*_0_)^1/2^, that is approximately ~4nm, i.e. the de Broglie length at room temperature. The time scale τ for exploration of consensus sequence is based on charge diffusion over spatial distance *R*=(2*Dτ*)^1/2^. By introducing the number of bases *N*=*R*/*a*_0_, where *a*_0_=0.34nm is the base-base distance, we obtain τ=(*Na*_0_)^2^/2*D*=*N*^2^/2*k*, where *k=(4π*^2^/*h*)|<*i*|*H*|*j*>|^2^*F* (*h* is the Planck’s constant, |*i*> and |*j*> denotes the initial and final states and *F* is the Franck-Condon density for the vibronic states) is the rate of multistep hopping dynamics of the π-electrons injected in the DNA base by the protein-DNA contact points. Basically, we suggest that initially base readout of protein recognizes dimers in the sequence to be identified. This suggestion is made necessary as in QW the starting position of the walker plays a fundamental role due to the CPE. For example, to identify a sequence 5’-GATATC-3’, there is an advantage to detect initially a dimer GA because in a primate genome of ~10^6^ bps (base-base distance 0.34nm), TA dimers appear approximately 28 thousand times, while GA approximately 100 thousand times [46–48]. Consequently, TA dimers are distant from each other on average about ~12-13nm, while this length is reduced to ~3.4-3.8nm for GA dimers, i.e., GA dimers distances fall in the de Broglie length for π-electrons at room temperature.

π-π entanglement is made possible if four energetic conditions are satisfied simultaneously. First, resonant coupling between the protein-DNA binding initial contact point and vibronic levels with vibronic manifold of the primary ion pair. Second, near degeneracy of the vibronic manifolds of the ion pair states located at the first nucleotide bases and the base preceding the acceptor state. Third, near degeneracy of the vibronic manifolds of the ion pair states located at the last nucleotide bases and the acceptor state [49]. In turn, the environment composed mainly by DNA vibrations expressed by thermally averaged nuclear Franck-Condon density, water mediated interactions, hydrogen fluctuations and ionic Na^+^, Mg^+^ stabilization operate as a long-time coherence crutch.

In summary, the proposed mechanism of Protein-DNA target searching problem, schematically represented in Figure 1, can be outlined as follows (limiting ourselves to consider as an example a typical sequence of endonucleases a 5’-GATATC-3’ sequence and protein base-readout regime):

**Figure 1.**
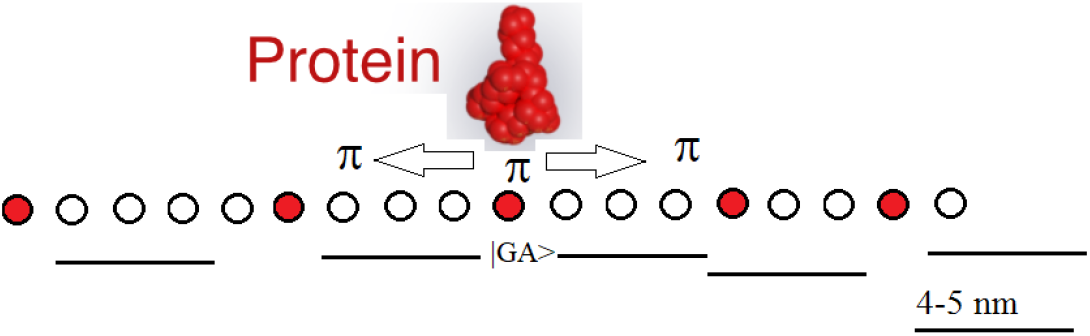
Schematic Sketch of QW applied to Protein-DNA binding recognition. The π-electron (walker) moves along the DNA chain by first identifying the GA dimers (red circles) locating a π electron and then completing the sequence hopping by GA dimer in GA dimer until the correct (consensus) sequence is identified by the other entangled π.

i. Any single DNA nucleotide is considered as a Hilbert state, where the nucleotides are organized in a tight-binding chain. The proteins does not distinguish between 3D or 1D dimensionality, and first detect a dimer of the sequence of interest, for example the initial dimer |*GA*〉. The contact is strongly conditioned by nucleotide-amino acids π-π interaction. One π is localized on |*GA*〉) and the other one (entangled to localized one) can walk through DNA chain.
ii. The π-electron performs a three-states QW on right or left to dimer traveling through HOMO-LUMO levels of DNA tight-binding sequence of bps. This speed up search, can identify the correct sequence for example by identifying another |*GA*〉 state, to which the protein jump and a new search starts, Figure 1.
iii. The walking π-electron shows a long-time survival probability and it is supported by the environment composed by thermal fluctuations, vibrational fluctuations of DNA backbone.

Furthermore, it should be noted that in QW the initial position plays a fundamental role, and therefore the initial dimer identified by the protein also defines the walk to be done, as will be highlighted by the asymmetry of the path described by the entangled π. It is well known that the total energy levels of HOMO-LUMO in the π-orbitals defines the hierarchical sequence of single bases, as C-A-T-G, and the various combination of dimers. Since the electronic properties of a DNA filament are dominated by π-electrons with one orbitals for site, the DNA can be represented as a tight-binding model with specific nearest neighbours interactions, along which the protein perform the QW.

In our application of QW to protein-DNA search problem, one key property is the memory of protein of exploration space of DNA. Memory is the ability of protein to take in account previously visited vertices in a self-avoiding mechanism [50]. Classical self-avoiding random walk has been already applied to protein folding [51]. In QW, self-avoiding mechanisms present a critical questions on how introduce memory update operator, able to update the protein memory as a function of the protein position along the DNA chains. This subject will be detailed in section 3.

## 2. Quantum and classical random walk picture of protein-DNA target search strategy

In our protein-DNA model, we consider a single protein molecule and a single DNA filament pictured as a chain of beads with various target sites. The DNA chain is supposed as having *N* discrete binds sites, and one of them at the position *i=1,…,N* is considered a target for the protein molecule. First, we wish to justify the tight-binding Hamiltonian model for the QW. Tight-binding Hamiltonians are commonly exploited to model coherent exciton energy transfer and thanks to the characteristics expressed in such transport problems can be closely related to QW systems [52–54]. In this section, we adapt the tight-binding Hamiltonian to the specificity, that is a strong prerogative of protein-DNA interactions, where the DNA nucleobases are free to engage with aromatic residues in π-π stacking, which interaction energies are estimated in water to range between −0.1 and −0.3eV [45]. In addition, we assume that DNA sequence prerogative is a first time problem, i.e., along a DNA chain, the protein search continues until the protein molecule reaches the specific sequence site for the first time [17,25].

Nonspecific protein-DNA adsorption energy can be divided into two major factors [55]. One is the sequence independent Coulomb energy of attraction between the positively charged domain of the protein surface and the negatively charged domain phosphate backbone. The other one is the sequence specific adsorption energy due to formation of hydrogen bonds of the protein with the DNA bases [56]. Position weight matrix (PWM) is a model that represents the specificity in tractable form, by probabilistic modelling or energetic modelling [57], by assuming that each position in the protein binding site contributes independently to recognition. Such model assigns a score to each possible base in each position, the sum of which gives a total score that allows an estimation of specificity.

During the searching dynamics, it is reasonable to assume that only sequences with binding energies in a certain interval are selected. If the binding domain of the regulatory protein is conserved throughout the searching dynamics the consensus sequence correspond to strongest binding sequence, hence any base-pair mismatch in the sequence will weaken the binding energy by a certain discrimination energy that depends on both the position and the mutated bps. If all positions are equally important and mutation contributes the same discrimination energy, then identifying the sequence energy is equivalent to specify the number of bp mismatches [58].

Since target recognition is often mediated by hydrogens bonds and the recognition α-helix interacts with several bps, many hydrogens may or many not contribute to specific or non-specific binding events. Recently, Tan and Takada proposed a coarse-grained model of the sequence-specific protein-DNA interactions [59]. In a coarse-gained model for a given complex structure or protein in binding relation with its recognition element, an amino acid in protein and a base in the DNA, are considered in contact when at least one of heavy-atoms of the *j*-th amino acids is within 0.34nm from one the atoms of the *m*-th base of the *B*_*m*_ bases. Inside the *m* base, the specific sequence is represented by an array of *m*′ bases, as for examples, GATATC, or GAATTC, for EcoRV, or EcoRI, respectively.

Based on relevant role of π-π stacking interaction between aromatic residues and nucleobases we utilize a simplified representation of coarse-grained model of protein-DNA interaction composed of three energy contributes. The quantum equivalent of such coarse-grained model is the tight-binding approach representing the random sequence of bps. In tight-binding picture, we need the onsite energies of bps and the hopping integrals between successive bps. The π-π interactions between stacked bp in double-strand DNA could provide a pathway for charge migration along DNA filaments, for example, when excess of charges localization at specific nucleic bases. The model of long-range electron transport is based on the scheme *d-B*_*n*_-*a*, where *d* plays the role of donor, and *a* the correspondent acceptor bases, *B*_*n*_ is the sequence of *n*-nucleobases. The electron transfer will occur for off-resonance coupling between the initial (doorway state) *d*\*{*B*_*n*_}*a* and low vibronic states (≅*k*_*B*_*T*=0.026eV) with all the vibronic manifolds connected to 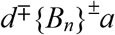. The electronic coupling can be expressed as (*γ*_d_*γ*_a_//Δ_1_)(*γ*/Δ)^*n*-1^ where the nearest-neighbour electronic matrix elements are 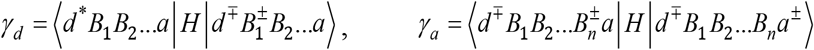 and 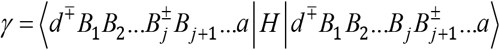, where *H* is the Hamiltonian of the system’s electronic and the states denote the valence bond electronic states. The energy gaps are Δ_1_=Δ*E*+δ and Δ=δ, respectively, where Δ*E~* few eV is the electronic energy gap, and δ~0.04eV is the reorganization energy. The rate of electron transfer along a specific sequence of length *L*=*N·a*_0_, where *N~150* is the number of bases and *a*_*0*_ is nearest-neighbour base-base distance, will be 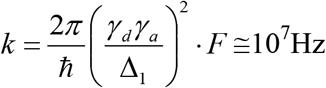 where, (*γ*_d_, *γ*_a_)~ 0.1-0.4 eV *F* ≅10^−5^ is the Franck-Condon density between the donor vibronic doorway state |*d*^*^*B*_1_*B*_2_…*a*〉 and the state 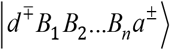. One long-range possible mechanism of electron transfer could be provided when resonant coupling between the electronic doorway state and (low) vibronic levels of the primary ion pair 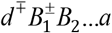, is associated to near degeneracy of states 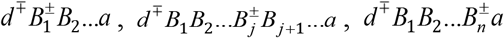 and, in turn, degeneracy of the vibronic manifolds of the base ion pair state 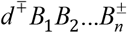 and the ion-pair state and the state 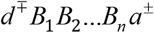.

In our model, we hypothesize that the electronic structure and carrier transfer travels through HOMOs and LUMOs levels, respectively. Therefore, we use the HOMO or LUMO onsite energies of bps and the HOMO or LUMO hopping integrals between successive bps. A tight-binding HOMO-LUMO Hamiltonian of a B-DNA sequence can be written as [60–62]

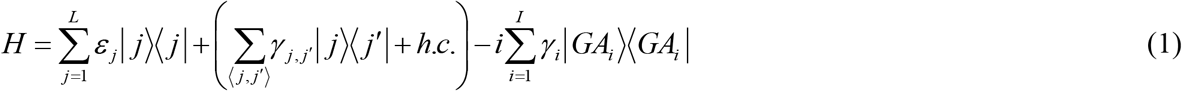

where *ε*_j_ is the HOMO or LUMO onsite energy of the *j-*th bp and 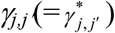 is the HOMO or LUMO hopping integral between the site *j* and *j′*, where <*j|j′*> = δ_j,j′_ indicates summation over all relevant neighbours, *h.c*. denoted Hermitian conjugate, in turn, the non-Hermitian contribution coincides with GA dimers. The non-hermiticity of Hamiltonian (1) due to the dimers of sequence to be identified by the proteins makes the eigenvalues complex introducing a term *iH*_*ii*_ in diagonal elements. The imaginary contribution to energy eigenvalues induces the localization to dimer states. In appendix B, we present a detailed description of non-Hermitian Hamiltonian for the protein walker.

The HOMO-LUMO representation of DNA sequence of length *L* is expressed as [61]

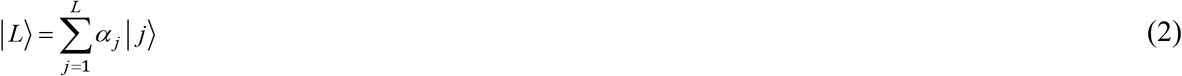

for example, a sequence target like 5’-GATATC-3’ is expressed as

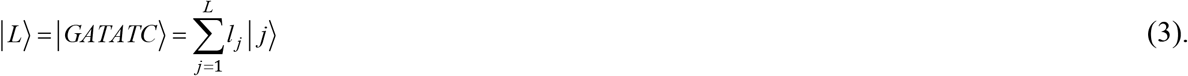

The time-independent Schrödinger equation

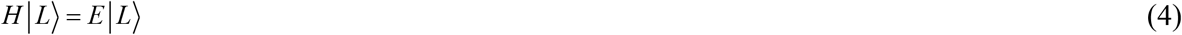

generates a system of *L* coupled equations of the form

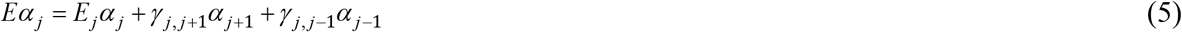

In the time-dependent the sequence *L* is

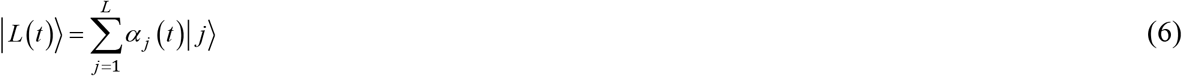

where |*α*_*j*_(*t*)|^2^ is the probability of finding the carrier at the *j-*th site at time *t*. The time-dependent Schrödinger equation

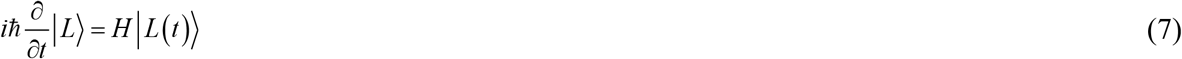

generates a system of *L* coupled equations of the form

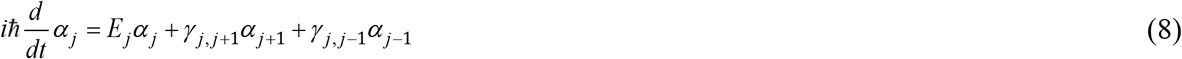

The *E*_*j*_ can be the on-site energy for the bps, ranging from approximately 8eV, for HOMO value of A-T or G-C, to 4.5-4.9eV for LUMO values [63], or the π-π* energy transition, 3.4-3.5eV for the two bps, respectively. Once the tight-binding matrix that is a symmetric tridiagonal matrix is solved evaluating the *L* eigenvalues, *E*_*k*_=*E*_*bp*_+2*γ*^bp^cos(2π*k*/*L*) and the correspondent eigenvectors, so that a quantity *x*(*t*) can be expressed as

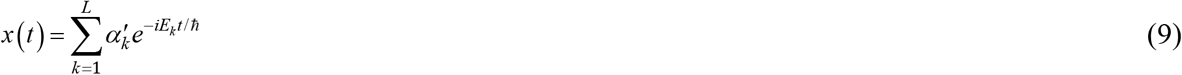

where the probability to find an electron at site *k* is now 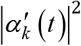.

In turn, there is another peculiar quantum feature in the protein-DNA target search. The ability of proteins to identify target on DNA chains is strictly connected to the efficiency to visit all sites of a given graph. This ability is commonly recognized as *cover time*. Cover time, that in quadratic speed up QW changes as *t*≅*N*^1/2^, refers to the capacity of exploration of a graph and not only to identify a specific target. A common mechanism to improve the efficiency of QW to cover all the sites of a graph is the self-avoidance ability [64], that requires that the protein remembers the site already visited and avoids positions where it has previously visited. Memory effects in quantum systems are commonly described by non-Markovian processes [65–66], however, continuous time QW are generally described by Markovian processes. Consequently, self-avoidance in continuous time QW could be obtained by introducing a complement operator composed by a sequence of qubits {*yes*=1, *not*=0} associated to site *i*. Unfortunately, this memory process is limited to relatively small evolution time because the size of Hilbert space grows exponentially with the size of the *N* graph vertices as *N*·2^*N*+1^. This problem can be overcome by invoking a non-Hermitian Hamiltonian *H*=*H*_0_-*iH*, where the imaginary term contributes for a correspondent imaginary term in the eigenvalues responsible for target localization.

The next section is focused to describe QW operating on restriction enzymes EcoRV and EcoRI.

## 3. 1-D searching strategy and coin-position entanglement. EcoRV and EcoRI as case study

In this section, we adopt the 1-Dimensional QW to describe the best-characterized endonucleases, the restriction enzymes EcoRV and EcoRI from *Escherichia coli*. Endonucleases are enzymes able to cleave the phosphodiester bonds within a polynucleotide chain. In particular, restriction endonucleases cleave only at high specific nucleotide sequences. More than 3000 type II restriction endonucleases have been currently identified, representing over 200 different sequence specificities [67]. The high specificity recognition of restriction enzymes together with their DNA scissoring features makes such enzymes important systems for understanding protein-DNA interactions. EcoRV consists of a dimer of 245 amino acid residues per monomer. Its enzymatic role is to cleave the foreign sequence 5’-GATATC-3’ at the initial GA step in a blunt-ended fashion, generating 5’-phosphate groups in a Mg^2+^-dependent reaction [68]. In the sequence, the initial GA is directly recognized only via hydrophobic contacts with the thymine methyl groups and the DNA is sharply bent by a 50° roll into a major groove at this position, unstacking the base. In a complex EcoRV-DNA, the DNA filament is located in a cleft between the two protein chains and is contacted to two peptide loops from each monomer together with other segments of the protein next to DNA phosphate groups. Analogously to cognate TA-sequence GATATC, a non-cognate AT-sequence, GAATTC is recognized by another restriction enzyme, EcoRI. The experimental evidence that the capacity of EcoRV to bend the AT sequence site is strongly limited to that of the cognate complex EcoRV-TA [55]. Analogously to EcoRV, EcoRI is a restriction enzyme in Escherichia coli able to recognize a 5’-GAATTC-3’ sequence, and cuts it, where now the central bp is composed by AT. EcoRI identifies GAATTC sequence over miscognate sites, normally one incorrect bp, by as much as 10^11^-fold [55]. When EcoRI binds to the specific DNA recognition site, two sort of arms protruding from the main domain of the endonuclease undergo to a phase of critical contacts to bases and phosphates in a flanking context on cognate site involving wild-type or A138T sequence, *in a so-called disorder-to-order transition* [56]. Hence, the protein-DNA contacts stabilize the specific complex consisting in the transition where the binding energy is utilized as a catalytic step. In this process, the complexes that deviate by two or more bps are completely discarded for cleavage.

Recently, Kurian et al. proposed a quantum mechanical model to clarify the mechanisms underlying type II endonucleases [45]. They proposed that type II endonucleases coordinate their two catalytic centers by entangling electrons in the target phosphodiester bonds through dipole-dipole couplings in the bound DNA sequence. In such model, collective electronic behaviour in DNA helix is characterized by quantized dipole-dipole oscillations through boundary conditions imposed by the enzyme. The collective electronic behaviour in DNA is simulated through interaction between delocalized electrons. These electrons belong to the planar-stacked bps that serve as *ladder rungs* stepping up the longitudinal helix axis. Where any rung of the helix can be represented as an electronically mobile sleeve vibrating with small perturbation around its fixed positively charged core due to Coulombic interactions between electrons clouds originated by induced dipole formation due to London dispersion from nearest neighbours. The enzyme acts as a decoherence shield making possible the quanta to be preserved in the presence of thermal noise by the enzyme’s displacement of water surrounding the DNA recognition sequence. One critical issues addressed by the model is that persistent correlations in DNA across longer spatial separations is a signature and persistence of entanglement at biological temperature.

Long-range electronic transport plays a leading role in CPE. In particular, the protein walker exploits long-range π-electron transfer to discriminate between nucleotides bases. Sequences *d-B*_*n*_-*a* can be identified by different hopping rates between bases. Measured hopping rates for A-A or G-G present values to be 1.2×10^9^ and 4.3×10^9^Hz, respectively [69]. Long-range electron transfer is based on multistep hopping mechanisms and intra- and inter-strand hopping rates, as well as activation energy, between nucleotides depends by the bases involved. For example, GAC presents an intra-stand hole hopping rate ~10^6^Hz and an activation energy ~0.3eV, while GAAC ~10^4^Hz and ~0.53eV, respectively [69]. Sequences as GAG and GAAG, again, present the more relevant changes due to insertion of an additional nucleotide A, going from ~5·10^7^(Hz) and ~0.2eV for GAG, to 10^5^(Hz) and ~0.45eV. Another sequence discriminant factor is the excess electron injection in DNA. The rates of electron injection depends by the bases sequences ranging from 10^12^Hz to 10^8^Hz. This is because any different nucleotide works with its specific electron acceptor and sequence specificity can modulate the electron injection [70]. Our supplementary assumption is that the protein exploits the electronic properties of DNA to speed up recognition of sequences by recombining the electronic properties in a CPE mechanism.

QW representation of such enzymatic recognition characteristics of GATATC sequence starts with the assumption that the DNA filament represents a 1-D system, Figure 1. The protein quantum walker can move only along two directions, right or left. The default initial position |0〉 is coincident with the double base |*GA*〉, i.e., |0〉 = |*GA*〉. Based on such initial condition, the quantum walker has three choices, to move right or left or to stay put. The Hamiltonian of such QW process is described by the relation H = H_DNA_ ⊗ H_C_ where H_DNA_, as tight-binding Hamiltonian in Eq. (1), represents the DNA filament, Eq. (2), including the Hilbert space which has orthonormal basis given by the DNA bases positioned at |*x*〉. H_C_ represents the coin operator and denotes a Hilbert space spanned by the orthonormal basis {|*r*〉, |*o*〉, |*l*〉}, where *r* stays for a right movement of the walker, *o* for stay put, and *l* stays for a left movement. The basis state is hence |*x*, *c*〉, where *x*∈DNA represents the position of the enzyme along the DNA filament, Figure 1, and *c*∈{|*r*〉, |*o*〉, |*l*〉} represents the coin state. The evolution of the walking enzyme at each step can be described by the global unitary operator, *U*, defined as *U* = *S* (*I* ⊗ *C*), where *S* is defined by *S* = |*x* − 1, *r*〉〈*x*, *r*| + |*x*, *o*〉〈*x*, *o*| + |*x*,*l*〉〈*x* + 1,*l*|, with suitable rate coefficients as in Eq. (1), and *I* is the identity matrix which operate in H_DNA_, while *C* is the coin operator. The *C* coin operator is built so that when the walker meets a state |A〉 on the left (palindrome sequence) of |*GA*〉 or a state |T〉 on the right so that the sequence identified can be 〈*A| 〉GA|* or |*GA*〉 |T〉, respectively. What is should be observed in such QW process is an asymmetry due to |*GA*〉 location in the sequence, as shown in Figure 2. In correspondence of such states the Coin operator is switched on |*x*_*A*,*T*_, *o*〉. This mechanism is very close to Lévy motion [71], which corresponds to random walk for most of the time spent around a state |*GA*〉, but when the sequence cannot be identified, the enzyme experiences large jumps, known as *Lévy flights*, to another dimer and hence actives another random walk around the new dimer. In turn, it should be remarked that only a three-states discrete QW simulate the recognition of a consensus sequence based on π-π entanglement represented in QW by the entanglement between position and coin [72]. The state of the walking enzyme, Ψ, is denoted by a tri-dimensional vector as

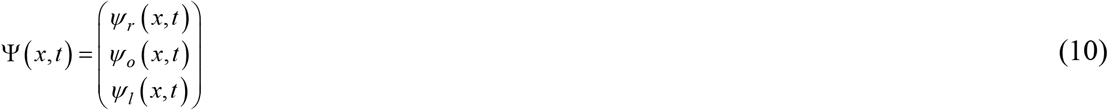

**Figure 2.**
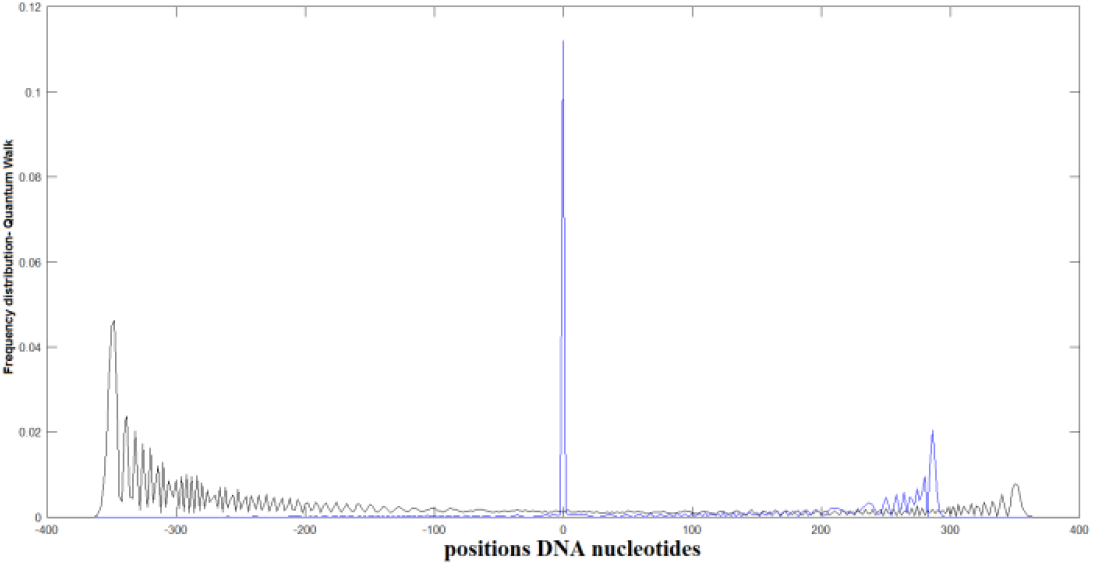
Three-state QW with initial state |0〉 = |*GA*〉, A, with CPE denoted by theta=1/3. Two-state QW comparison to standard QW without CPE. Due to specific sequence to be identified, both the QWs are asymmetric.

The walker evolves under the shift operator as

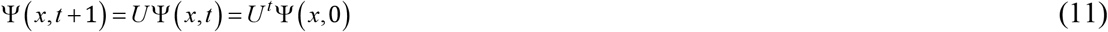

where *U*=*S·C* is the global unitary operator combination of the Shift operator with Coin operator. In momentum space, the relation (11) can be written as 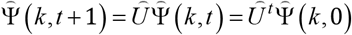, where the inverse transform is

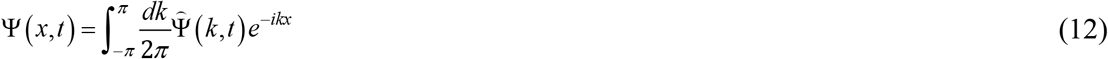

where the global unitary operator 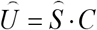 now the Shift operator takes the simple form [73]

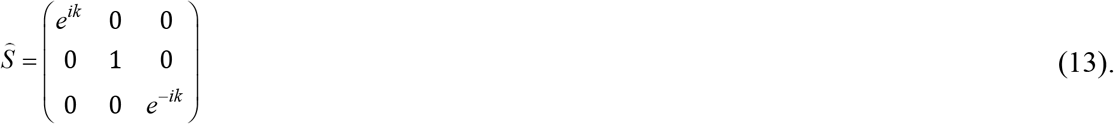

As Coin operator, we have different possibilities, here, we adopt the following operator [73]

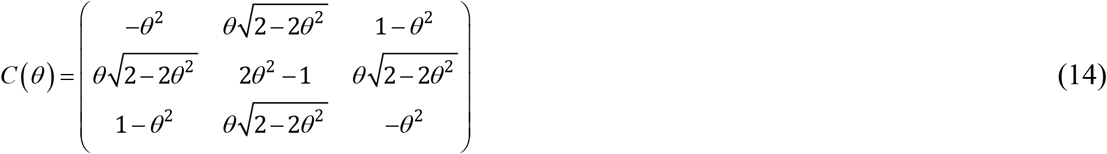

where the coin operator θ∈(0,1) and for any of such value the rows are mutually orthogonal and for 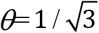 it is equal to the Grover operator. A Coin operator (14) possesses the fundamental property that one of the eigenvalues is 1, generating the rest state that in protein search corresponds to |*GA*〉 state. Consequently, such Coin operator simulates the enzyme strategy to fix an initial state and to taste possible subsequent nucleotides, if the local search does not give the wanted results, the enzyme can shift along the DNA chain and identify another dimer and so go on. In Figure 2, we report the probability distribution of the three-states QW (blue line) with CPE and compared to normal QW (black line). Three-states search strategy that provides the localization of a dimer has the disadvantage to limit the quantum ballistic property, as evidenced in Figure 2, where the total positions investigated by the QW with a localized state (blue line) are less than a two-states quantum walker that does not localize a specific state and hence the total position spanned is greater (black line).

The fundamental role of CPE in the search of a sequence starting form an initial state grounded on a central dimer |*GA*〉 can be explained as follows. The entanglement measure is a parameter usually used to quantify the incidence of entanglement between two subsystems, in our case, the position space of EcoRV and the coin space. The entanglement *E* can be evaluated making use of von Neumann entropy *S*(*t*), known as the entropy entanglement [74]

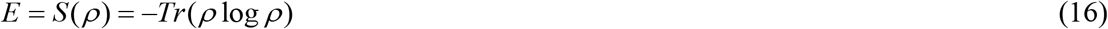

where *ρ* = |Ψ〉〈Ψ| is the density matrix, that should be calculated on a bipartite system composed by two subsystems, position space and coin space, respectively. The maximum value of the entanglement *E* between the two subsystems can be expressed as *E*_*max*_=*log*_2_(*min*(*d*_1_,*d*_2_)), where *d*_1_ and *d*_2_ are the dimensions of the two subsystems. For a normal one-dimensional QW, *E*_*max*_=*log*_2_(2)=1, while for a three-state QW *E*_*max*_=*log*_2_(3)≅1.58. This is because the dimensions of the coin space in a three-state QW is bigger than one-dimensional correspondent standard QW involving a higher entanglement between position and coin. It is remarkable that third state for coin operator, other than right or left states, increase the entanglement [73]. Another feature of CPE of the protein walker in specificity recognition need to be emphasized. In discrete QW, CPE plays a leading role in maintaining long-time coherent superposition features between coin and position resistant to environmental dephasing role due to water molecules fluctuations and thermal vibration. The CPE has shown that small amount of decoherence helps the speed up behaviour of QW [75], and maximally entangled coins can be obtained by the existence of rest sites that are position states that allows, when applied to biology, the protein walker to stay at a current vertex while exploring the surrounding positions.

## 4. Discussion

Common stochastic modelization applied to protein-DNA search target problem provides a DNA molecule represented as a chain consisting of *L*+1 binding sites, one of them being the target site to be identified by the protein [17, 76–77]. In such picture, the protein starts to search from the solution, and it diffuses very fast in the volume solution around the DNA reaching to all segments of the DNA with equal probability, with a binding rate *k*_on_. Once bound, the protein can slide along the DNA chain with a diffusion rate *u* with equal probability in both directions with a classical random walk and can dissociate from DNA with rate *k*_off_. Despite the simplicity of such modelization, some aspects seem to be clear an universal in the protein-DNA interaction, other remains still unclear, for example the experimentally evidenced short time (seconds) to identify consensus sequence in enzyme restrictions, and how noncovalent forces can operate at room temperature with high rate of efficiency despite in presence of not negligible environmental fluctuations. A more recent different theoretical approach, within the family of facilitated diffusion, takes into account the correlations between spatial motion (one-dimensional and three-dimensional) and the non-specific interactions [77]. One feature of such approach is the mean first-passage time for one-cycle searching and the total search time is simply τ=(*L/λn*_ads_)τ_1_, where *L* is the DNA length, *λ* is the target length, *n*_ads_ is the number of proteins adsorbed on the DNA under investigation and τ_1_ is the search time for one cycle. One prediction of the model that is partially confirmed by some experiments is that the protein spends most of the time on DNA, and that non-specific interactions play a leading role in facilitated diffusion. This is because although non-specific binding proteins move slower along the DNA slowing down the search, however, non-specific interactions stimulate more protein to bind to DNA and searching in parallel accelerating the search. One critical aspect of correlation model is the assumption that search time takes place faster than the relaxation to equilibrium for binding to DNA. To clarify such two critical questions, we have introduced a quantum version of CRW. This need some conceptual changes to the protein-DNA search target problem. The DNA chain is composed by Hilbert states, the first binding state on DNA cannot identified by the protein based on equal probability, but the first binding state is a dimer state part of the sequence to be recognized. Hence, from the starting states, the protein exploits the CPE to search the consensus sequence, via a speedup donor-acceptor mechanism involving π-electrons moving quadratically along the DNA segments, able to scan the DNA nucleotides for distances roughly equivalent de Broglie length. This macroscopic quantum effect occurs thanks to decoherence shield of environment (hydrogen bonds, thermal and vibronic DNA backbone states) protecting the superposition of states during the QW process. Mohseni and Aspuru-Guzik have demonstrated that continuous time QW in photosynthetic energy transfer is environment-assisted and while an unitary Grover search mechanism does not match the high efficiency of photosynthetic process, this efficiency can be achieved by open nature of the photosynthetic energy transfer with fluctuations in protein and solvent [78]. The environment plays a role to improve the efficiency and this role is described by a non-hermitian contribution generated by the coupling of the system of interest with environment [79].

Another advantage of quantum approach is to bypass the delicate question of the two timescales, fast for the first binding in a 3D context, and slow for the 1D refinement of the search for the complete sequence. Currently, stochastic models based on facilitated diffusion have combined such two time scales with *lowering of dimensionality* [25]. The concept of *lowering of dimensionality*, proposed initially by Berg, Winter and von Hippel (BHH) [80], provides a search process based on a consequence of events the include a binding to non-specifically DNA site form a 3D space. Hence, a 1D movement along the DNA contour to identify correct sequence with possible unbind if no target is found and binding to another site and so gone. However, such model cannot explain the fast association rates for the protein search on DNA because it provides similar values for 1D and 3D diffusion value, contrarily to what demonstrated experimentally. This is because, the protein-DNA search strategy is based on an interplay of different spatial dimensionality, essentially between 1D chain sliding and 3D features. The fast association rates on 1D and 3D originate from strong electrostatic attractions between searching protein and DNA sequences, as well as from reduction of dimensionality in the target searching strategies. The hypothesis of leading mechanisms implying reduction of dimensionality is at the base of facilitated diffusion that provides the lowering spatial dimensionality by combining 1D diffusion along the DNA chain contour (sliding) with 3D DNA segment dissociation and consequent jump to another DNA filament. The 3D mechanisms are essentially a combination of dissociation-reassociation events of short life spans, hopping events, intersegmental jumps between nearby DNA segments and larger volume 3D diffusions [81–82].

The formulation of first-passage-time picture applied to identification of DNA targets by Kolomeisky *et al*. is independent of the initial starting position of the searching enzyme where an initial uniform distribution of enzyme positions is commonly assumed [21]. The first-passage-time distribution arises from survival probability distribution. Survival probability for the entangled walking π-electron moving through a QW process along a tight-binding Hamiltonian is represented in Figure 3 and compared to the equivalent classical quantity, see Appendix B. Figure 3 shows that quantum survival probability for the π-electron displays a long-time behaviour, which has no counterparts in the classic case. Since decoherence can play a role in breaking the CPE, the tight-binding Hamiltonian is formulated to take in account the effect of environment by introducing a non-Hermitian term.

**Figure 3.**
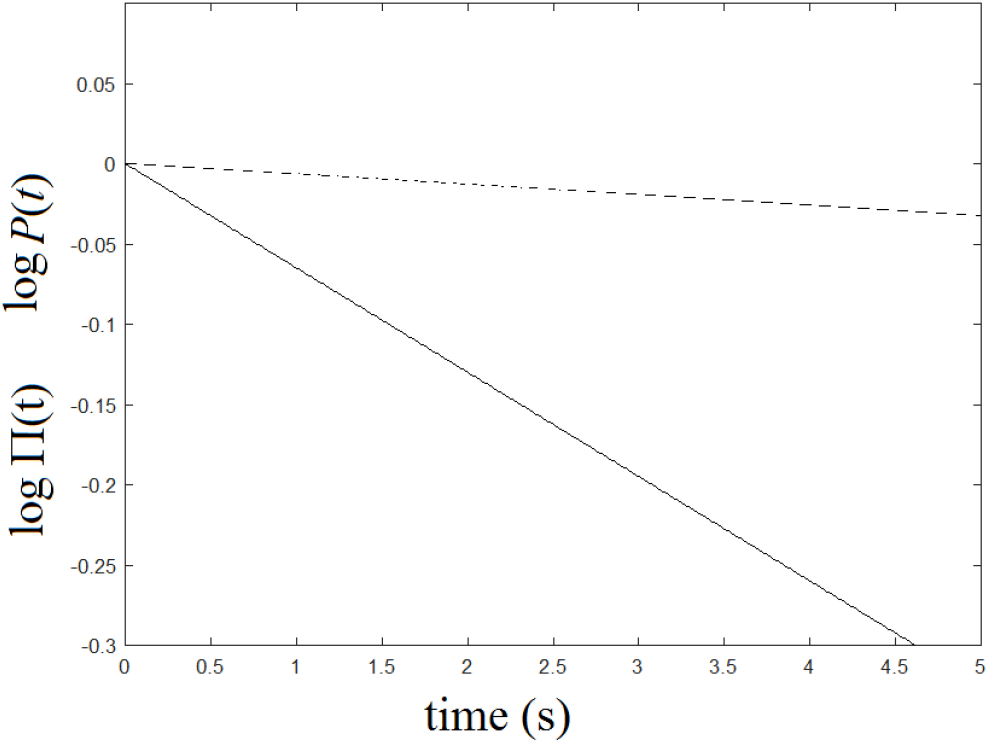
Temporal decay of classical survival probability *P*(*t*) (dashed line) and the quantum survival probability *Π*(t) (solid line) for *N*=150 nucleotides in semilogarithmic scales.

The persistence of CPE even under the effects of environmental fluctuations can be explained as follows. Let us consider the density operator formalism to describe the nonunitary evolution of the discrete QW [83]

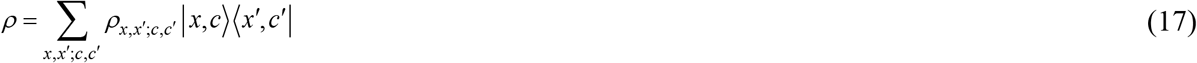

where the matrix *ρ*_*x,x′;c,c′*_ denotes the probabilities of occupying some given nucleotides and coin states (diagonal terms) and the amount of coherence between quantum states |*x*, *c*〉 and |*x*′, *c*′〉 corresponding to off diagonal terms. If we assume decoherence process takes place at the end of any step so that the time-scale of decoherence is comparable to time-scale of quantum walker, on each step, coin decoherence leaves the coin-diagonal elements unchanged. On the contrary, it reduces the coin-off-diagonal elements, *ρ*_*x,x′;c≠c′*_, by a factor 1-*η*_*c*_. Analogously, spatial decoherence leaves unchanged position-diagonal elements, and it reduces on each step the position-off-diagonal elements *ρ*_*x≠x′c,c′*_ by a factor 1-*η*_s_. This description of decoherence takes in account the possible role of thermal noise on hydrogen bonds between nucleotides. Under decoherence the asymptotical behaviour of the quantum walker the variance of the diffusive spreading can be expressed by 〈*x*^2^〉 ≅ *Dn*. For a process of walk combining quantum and CRW behaviour, the diffusion constant can be expressed *D* = [1 + (1 − *η*_*c*_)^2^]/[1 − (1 − *η*_*c*_)^2^], where *η*_c_=0 denotes a totally coherent process, and a divergent diffusion constant, and a decoherent process, *η*_c_=1, where the diffusion constant assumes the traditional classical constant value. This implies that during the QW process the walker manages the coherence-decoherence transition according to the needs of the research path, remaining in an intermediate degree between coherence and decoherence. Analogous results can be obtained in another bipartite system, if we consider bipartition between sections of DNA chains arounf the initial dimer. A bipartition that provides a subsystem A=[-*L*/2, *p*] of nucleotides around GA dimer, and the rest of the DNA chain as the subsystem *B*=[*p*, *L*/2] composed by the nucleotides which remain at the next dimer, which relative distance is *L*, and *p* is the current position of the walking π-electron, and the von Newmann entropy is calculated making use of the reduce density matrix *ρ*_A_(*t*)=Tr_B_*ρ*(*t*).

In turn, one interesting exploitation of protein-DNA searching mechanism is the possibility to build new type of computers. Quantum computing is limited by the dissipation of coherence due to the coupling with environment. It has been conjectured that biological systems exhibiting room temperature quantum coherence such light harvesting complexes could provide new computational devices realized with different component basis achieving new magnitude scales for miniaturization and speed. In a computational framework we have usually to find the minimum of some complex functions that in many cases can take discrete values. Recently, continuous-time QW has been employed by Fahri *et al*. to simulate a NAND (NOT-AND) component and recent experiments tried to realize it on molecular systems [84]. A NAND gate presents a total of *N* inputs but only a single output, making possible the computation of arbitrary Boolean function of the form *F*:{0,1}^*N*^→{0,1}. If protein-DNA binding search is based on QW, as proposed in this paper, than such biosystem could simulate a NAND-tree. In the idea of Fahri *et al.*, a NAND-tree is a binary tree of NAND gates containing *N* inputs and a single output. The time-independent Hamiltonian of the entire system is simply *H*=*J·G*, where *J* is the coupling constant and *G* is adjacent matrix. The input ***x*** is encoded by the on/off coupling of the input nodes. An elegant approach to NAND-tree has been recently proposed by Wang *et al*. [85]. They propose to attach to the runway a sort of quantum slide, i.e., a chain with a parabolic shape able to scan the site-to-site coupling profile. A single photon or a light pulse, representing a Gaussian wave-packet, is injected into a single site located at the edge of the quantum slide generating the QW. It is immediate that protein-DNA system can represent an analogous large scale system of such NAND-tree, where if the protein recognizes a component basis of the correct sequence, the computation outcome is *F*(***x***)=1, or, on the contrary, *F*(***x***)=0. By measuring whether the protein has recognized dimers of sequence components or not, the binary computation outcome can be established by measuring the distribution of recognized sequence basis versus the evolution distance travelled by the protein to verify the NAND tree logic.

## Conclusions

Protein-DNA interactions are fundamental for life. Such interactions include, for example, binding selection to DNA, which is classified as: i) *Enzymatic*, where the protein modifies DNA; ii) *Transcription Factor* (TF), where genetic information contained in the DNA nucleobases is transcribed into RNA and new proteins are generated, or, iii) *Structural-DNA Binding Proteins*, where the protein supports DNA structure, DNA bending or to aggregate other proteins. Since the pioneering experiments by Riggs *et al*., in the early ’70 of past century, it has been increasingly demonstrated the extraordinary ability of proteins to find the targets on the DNA chains very quickly, in some cases, with two orders of magnitude faster than the diffusion limit. Although several theoretical mechanisms have been suggested to describe the origin of such impressive diffusional ability, however, none of such models currently has the ability to fully describe the protein search strategy.

Focused on the example of recognition of consensus sequences by EcoRV or EcoRI, we have suggested and introduced a mechanism based on Quantum Walk exploiting π-π entanglement in protein-DNA complexes. The Quantum Walk mechanism, that is the quantum version of classical Random Walk, provides that the walker (the protein) resides at a vertex in a graph (the DNA structural topology), locates, initially, a π-electron on a dimer of a sequence to be recognized and uses the other entangled π-electron to scan the DNA chain. Differently to classical Random Walk, where the walker moves randomly, the quantum walker spreads quadratically faster than classical Random Walk the different positions moving coherently on a superposition of different vertices. In addition, initial positions play a fundamental role in Quantum Walk, so enabling for the walker the possibility to distinguish between initial dimers, as evidenced by asymmetric walks. When applied to protein-DNA target search problem, Quantum Walk gives the possibility to achieve the experimental diffusional motion of proteins over diffusion classical limits experienced along DNA chains exploiting quantum features such as coin-position entanglement and long-time survival probability induced by environment. In turn, our results show that, under quantum picture, the protein search strategy does not distinguish between one-dimensional and three-dimensional case.

## Appendix A

Although, of intrinsically discrete nature of molecular interactions of protein-DNA complexes, the searching process can be considered continuous in time. For this reason, a CRW model based on continuous time is physically reasonable. Another advantage of a protein-DNA searching process based on continuous-time CRW is the possibility to describe different regimes, sliding- and jumping-regime, in a unique formalism. In turn, a continuous-time CRW is mathematically formulated to yield a quantum-mechanical Hamiltonian of tight-binding type since the formalism exploits the mathematical analogise between time-evolution operators in classical and quantum mechanics [60, 86]. As in CRW, QW can show a discrete or a continuous-time mode. QWs take place on a graph denoted by a pair {*V,E*} consisting of a nonempty, countable set of points *V*, representing the number of *N* sites (the DNA bases) making up the graph, joined pairwise by a set of links *E*. Discrete time QW is a bipartite system including Position and Coin operator, respectively, with Hilbert space *H*=*H*_*p*_⊗*H*_*c*_, where the position degrees of freedom specifies the location (vertex) of the walker in the graph and the coin degrees of freedom specify the direction the walker on which it proceeds [87]. Continuous time QW, in contrast to discrete QW, evolves under a Hamiltonian, which is defined with respect to the graph simulating the system under investigation [88]. In addition, no Coin operator is required. Another computational advantage is that continuous time QW leads on fairly straightforwardly from continuous time CRW. Under such assumptions, continuous time QW would fit better to match the protein-DNA interaction, because it depends on the underlying structure, since continuous time QW features strongly is affected by the interplay between the walk dynamics and the topology of the protein-DNA structure.

The quantification of speedup behaviour of variance by using continuous time QW on the graph simulating the DNA structure is made through the use of an adjacency matrix [89]

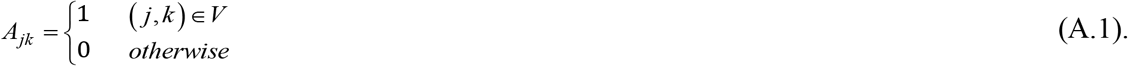

By denoting the transfer matrix of the graph with *N* bases as **T**, the conditional probability of the walker located the base *k*, at time *t*, when is started from the base *j* at time *t*=0 is [86]

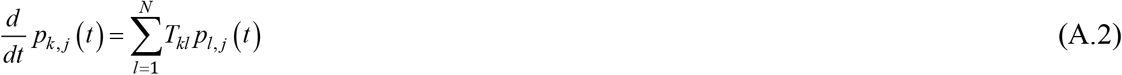

The quantum counterpart of Eq. (A.2) is obtained considering the Hamiltonian *H*=-**T** of the structure and the states |*j*〉 representing the walk at site *j* denote the whole Hilbert space and provide an orthonormal basis set, with the notation 〈*k|j*〉 = *δ*_*k,j*_ and 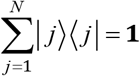. The wave vector can be written as 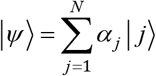, with coefficients given by α_j_ = 〈*j*|*Ψ*〉. Consequently, the analogous of Eq. (A.2) is the following Schrödinger equation

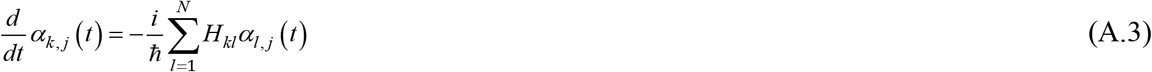

where α_*k,j*_ = 〈*k*|*e*^−*iHt/ħ*^|*j*〉 and the analogous quantum-mechanical transition probability is now π_*k,j*_(*t*) = |α_*k,j*_(*t*)|^2^. For the random walk, classical or quantum, the average displacements can be written, invoking the solutions of Eqs. (A.2) and (A.3), are 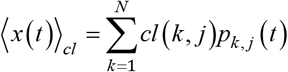 or 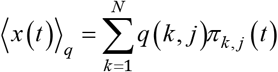, where subscript *cl* or *q* denotes classical or quantum, respectively. In quantum case, an additional average on all the starting points should be generally accounted. Finite systems present peculiar properties due to specific structure topologies. As a simplified example, we can consider the DNA as an infinite chain composed by *N* nodes, whose quantum amplitudes are

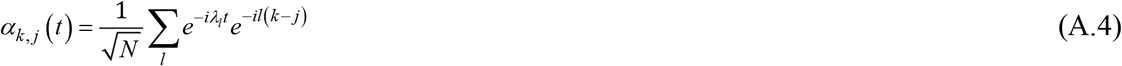

where *λ*_*l*_ is the *l*^th^ eigenvalue associated with the chain nodes. For high limit of *N* (condition satisfied by a genome) and substituting the sum over *l* by the correspondent integral, we obtain. *α*_*k,j*_(*t*)≅*i*^*k−j*^*e*^*−it*^*J*_*k−j*_(*t*). The calculation of transition probability amplitudes implies 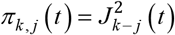. The squared Bessel function 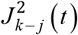 is almost everywhere positive. By exploiting the Bessel function properties [90], the average displacement in quantum case can be written as

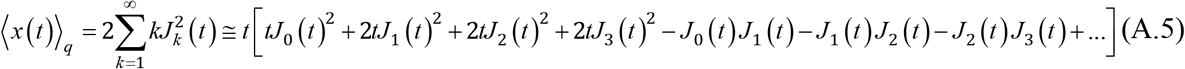

For long time the average displacement on an infinite chains reduces to<*x*(*t*)>_*q*_~*t*, this is because the asymptotic expansion of the Bessel function is

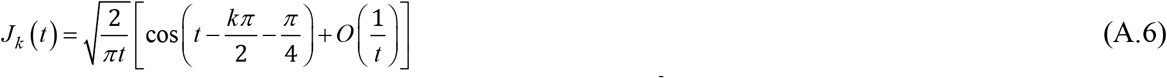

In figure A.1, we plot the average displacements <*x*^2^(*t*)> travelled by the protein along a DNA filament in classical and quantum case, for small and intermediate times.

**Figure A.1.**
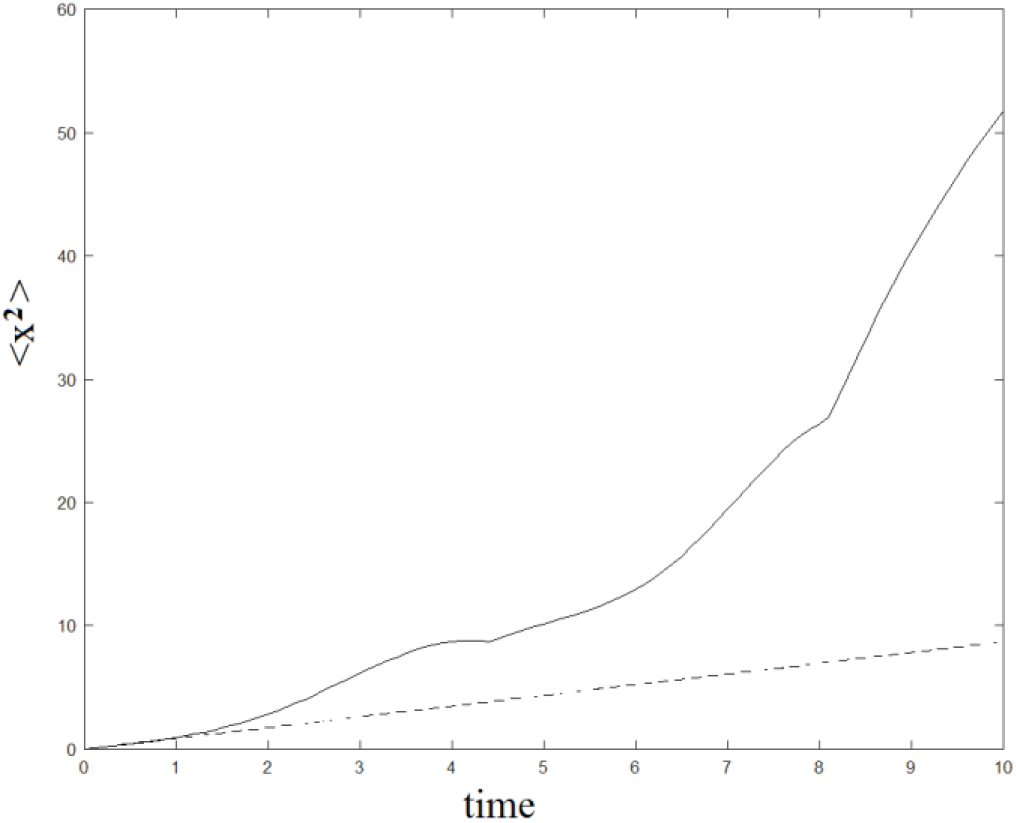
Quadratic speed up behaviour of variance, under QW picture (continue line) or CRW picture (dashed line).

## Appendix B

In order to quantify the dissipation role and decoherence process for the π-electron walking along the DNA chain, we adopt again the continuous time QW picture described by the following non-Hermitian Hamiltonian for the walker along the DNA filament, [60, 77]:

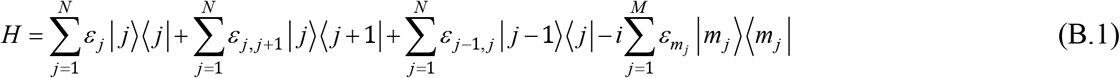

where *M* denotes the ensemble of the environmental channels denoting the thermal phonon bath connected to DNA chain, *ε*_j_ denotes the nucleotides energies, *ε*_*j,j±1*_ denote hopping integrals along the DNA chain, the coefficient ε_*mj*_ denote the strength of binding to the specific DNA site and N is the number of nucleotides located between two |*GA*〉 dimers. The cardinality of *M* denotes the strength coupling between protein-DNA system and environment. The second and third term on the right side of equation (B.1) are responsible for the states *r* and *s*, corresponding to sliding motion of the protein along the DNA chain. Due to the non-hermiticity of Hamiltonian (B.1), its eigenvalues are complex and can be written as *E*_*l*_=*ε*_*l*_−*iγ*_*l*_ (*l*=1,…,*N*), while the set of eigenvectors |*ξ*_*l*_〉 satisfy the completeness relation 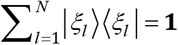. Due to complex eigenvalues, the dissipative role of environment in tight-binding model is analogues to traps located along the DNA chain. The transition amplitude *α*_*k,j*_(*t*) from the site *j* to the site *k* which obeys the following Schrödinger equation (A.3), which solution, given the Hamiltonian (B.1), can be written as

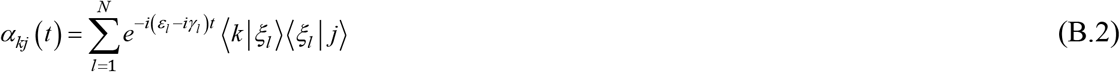

The sliding time of protein along the DNA chain can be calculated as follows. Let us consider *m*∈*M* denoting the targets on the DNA chain to be identified by the protein, we can exploit the relation 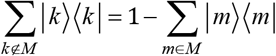 to write the sliding state as

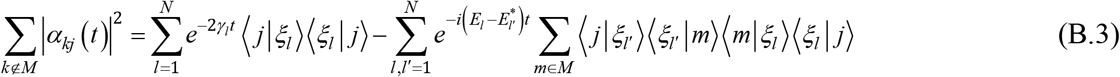

And, by averaging over all *j*∉M, the survival probability 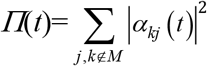, that coincides with mean sliding probability, becomes [91]

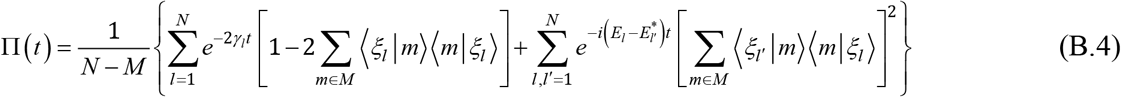

Note that two contributions are present, a decaying and an oscillatory term, while the classical counter part of Eq. (B.4) can be written [86]

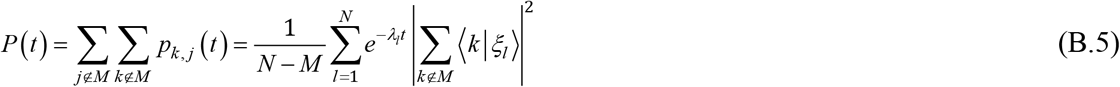

Temporal decay of Eq. (B.5) is defined by *γ*_*l*_, i.e. the imaginary contributions of *E*_*l*_. At intermediate and long times and for *M*<<*N*, Eq. (B.5) can be approximated by 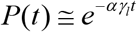 where *α* takes into account the ratio between the nucleotides visited in relation to the number of nucleotides present between two GA dimers. Eqs.(B.4 and B.5) describe also the survival probability of a π-electron moving between two GA dimers. Survival probability is strictly connected to cover time. Cover time takes an important role in random walk, because the probability to find a target is inversely proportional to cover time. Adopting the first-passage approach and self-avoiding mechanisms, the mean time required to cover all the N nodes in a given graph can be expressed in an one-dimensional case as <*t*_*cov*_>~*N*^*α*^, <*t*_*cov*_>~*N*(log*N*)^*α*^ in the bidimensional case [92], while in the three-dimensional case, we have <*t*_*cov*_>~*N*(log*N*) where α=1 in a self-avoiding picture and α=2 in classical random walk [92]. In the quantum case, the calculation of mean time to cover all the *N* nodes in the first-passage problem is more complex. As demonstrated by Childs and Goldstone, the spectrum of the Hamiltonian *H* can be understood in terms of the spectrum of the periodic or quasi-periodic structure. Hence, the quantum searching probability Π can be defined as the inverse of cover time in the quantum walk case, for a graph with *N* nodes and M targets in the following way [93]:

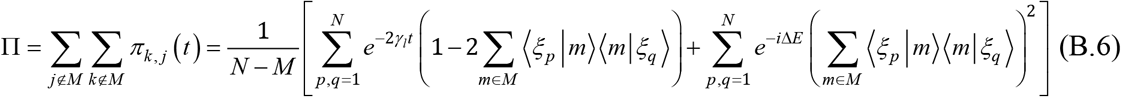

where Δ*E*=*E*_*p*_−*E*_*q*_. Plotting Eq.(B.5) and Eq.(B.6) we obtain the survival probability as displayed by Figure 3. It is evidenced that in quantum case presents long time for survival probability of π-electrons that which has no counterparts in the classic case [91].

## Acknowledgments

The author wishes to thank Patrizia Cioni, Edi Gabellieri and Eli Barkai for illuminating discussions and suggestions.

## References

1. Alberts, B., Bray, D., and Lewis, J. 2002. Molecular Biology of the Cell, 4^th^ ed., Taylor and Francis.

2. Rice, P. A., and Correll, C. C. 2008. Protein-Nucleic Acid Interactions: Structural Biology, The Royal Society of Chemistry, Cambridge, UK.

3. Redding, S., and Greene, E. C. 2013. How do proteins locate specific targets in DNA? Chemical Physics Letters, 570:1–11.

4. Rohs, R., Jin X., West, S. M., Joshi, R., Honig, B., Mann, R.S. 2010, Origins of specificity in protein-DNA recognition, Annu. Rev Biochem., 79:233–269.

5. Wilson, K.A., Kellie, J.L., Wetmore, S.D., 2014, DNA-protein π-interactions in nature: abundance, structure, composition and strength of contacts between aromatic amino acids and DNA nucleobases or deoxyribose sugar, Nuclei Acids Research, 42:6726–6741.

6. Gowers, D. M., Wilson, G. G., and Halford, S. E. 2005, Measurements to the contributions of 1D and 3D patways to the translocation of a protein alonf DNA, Proc. Natl. Acad. Sci. USA, 102: 15883–15888.

7. Blainey, P. C., Luo, G., Kou, S. C., Mangel, W. F., Verdine, G. L., Bagchi, B., and Xie, X.S. 2009, Nonspecifically bound proteins spin while diffusing along DNA, Nat. Struct. Mol. Biol., 16:1224–1229.

8. Halford, S. E. 2009. An end to 40 years of mistakes in DNA-rpotein association kinetics? Biochem. Soc. Trans. 37:343–348.

9. Barkley, M. D., 1981, Salt dependence of the kinetics of the lac repressor-operator interaction: role of nonoperator deoxyribonucleic acid (DNA) in the association reaction, Biochemistry 20:3833–3842.

10. Winter, R.B., Berg, O.G., von Hippel, P.H., 1981, Diffusion-driven mechanism of protein translocation on nucleic acids. 3. The Escherichia coli lac repressor-operator interaction: kinetic measurements and conclusions, Biochemistry, 20, 6961–6977.

11. Stanford, N.P., Szczelkun, J. F., Marko, J. F., and Halford, S.E. 2000. One- and three-dimensional pathways for proteins to reach specific DNA sites, EMBO J., 19:6546–6557.

12. Halford, S., and Marko, J. 2004. How do site-specific DNA-binding proteins find their targets? Nucleic Acids Res., 32: 3040–3052.

13. Riggs, A. D., Bourgeois, S., and Cohn, M. 1970. The lac repressor-operator interaction. III. Kinetic studies. J. Mol. Biol. 53:401–417.

14. von Hippel, P.H., and Berg, O. G., 1989. Facilitated target location in biological systems, J. Biol. Chem. 264:675–678.

15. Gilmore, J. L., Suzuki, Y., Tamulaitis, G., Siksnys, V., Takeyasu, K., Lyubchenko, Y. L. 2009. Single-molecule dynamics of the DNA-EcoRI protein complexes revealed with high-speed atomic force microscopy, Biochemistry, 48:10492–10498.

16. Tafvizi, A., Huang, F., Fersht, A., Mirny, L. A., van Oijen, A. 2011. M. A single molecule characterization of p53 search on DNA, Proc. Natl. Acad. Sci. USA, 108: 563–568.

17. Kochugaeva, M. P., Berezhkovskii, A. M., Kolomeisky, A.B. 2017. Optimal Length of Conformational Transition Region in Protein Search for targets on DNA, J. Phys. Chem. Lett., 8:4049–4054.

18. Gorman, J., and Greene, E. C. 2008. Nat. Struct. Mol. Biol. 15:768–774.

19. Loverdo, C., Bénichou, O., Voituriez, R., Biebricher, A., Bonnet, I., and Desbiolles, P. 2009. Quantifying hopping and jumping in Facilitated Diffusion of DNA-Binding Proteins, Phys. Rev. Lett., 102:188101.

20. Bénoichou, O., Chevalier, C., Meyer, B., and Voituriez, R. 2011, Facilitated Diffusion of Proteins on Chromatin, Phys. Rev. Lett. 106, 038102.

21. Shvets, A., Kochugaeva, M. P., Kolomeisky, A. B. 2018. Mechanisms of Protein Search for Targets on DNA: Theoretical Insights, Molecules, 23:2106.

22. Slutsky, M., Mirny, L., 2004, Kinetics of Protein-DNA interaction: Facilitated Target Location in Sequence-Dependent Potential, Biophysical J. 87, 4021–4035.

23. Cherstvy, A.G., Kolomeisky, A.B., Kornyshev, A.A., 2008, Protein-DNA Interactions: reaching and Recognizing the Targets, J. Phys. Chem. B, 112, 4741–4750.

24. Kolomeisky A.B., 2011, Physics of protein-DNA interactions: mechanisms of facilitated target search, Phys. Chem. Chem. Phys. 13, 2088–2095.

25. Kochugaeva, M.P., Berezhkovskii, A.M., Kolomeisky, A. B., 2017, Optimal Length of Conformational Transition Region in Protein Search for Targets on DNA, J. Phys. Chem. Lett., 8, 4049–4054.

26. Leven, I., Levy, Y. 2019, Quantufying the two-state facilitated diffusion model of protein-DNA interactions, Nucleic Acids Research, 47, 5530–5538.

27. Bonnet, I., et al., 2008, Sliding and jumping of single EcoRV restriction enzymes on non-cognate DNA, Nucleic Acid Research, 36, 4118–4127.

28. Monico, C., Capitanio, M., Belcastro, G., Vanzi, F., Pavone, F.S., 2013, Optical Methods to Study Protein-DNA Interactions *in Vitro* and in Living Cells at the Single-Molecule Level, Int. J. Mol. Sci., 14, 3961–3992.

29. Kamagata, K., Itoh, Y., Subekti, D.R.G., 2020, How p53 Molecules Solve the Target DNA Search Problem. A review, Int. J. Mol. Sci. 21, 1031.

30. Riley, K.E., Hobza, P., 2011, Noncovalent interactions in biochemistry, WIREs computational Molecular Science, 1, 3-17; Diaz de la Rosa, M.A., Koslover, E. F., Mulligan, P. J., Spakowitz, A.J., 2010, Dynamic strategies for target-site search by DNA-binding proteins, Biophysical J., 98, 2943–2953.

31. Mohseni, M., Omar, Y, Engel, G.S., and Plenio, M.B. 2013. Quantum effects in biology. Cambridge University Press, Cambridge UK.

32. McFadden, J., and Al-Khalili, J., 2018. The origins of quantum Biology, Proc. R. Soc. A 474: 20180674.

33. Marais, A., Adams, B., Ringsmuth, A. K., Ferretti, M., Gruber, J. M., Hendrikx, R., Schuld M., Smith, S.L., Sinayskiy, I., Kruger, T. P.J., Petruccione, F., and van Grondelle, R. 2018. The future of quantum biology, J. R. Soc. Interface, 15:20180640.

34. Kohen, A., Cannio, R., Bartolucci, S., and Klinman, J.P., 1999. Enzyme dynamics and hydrogen tunneling in a thermophilic alcohol dehydrogenase, Nature, 399: 496–499.

35. Page, C. C., Moser, C. C., Chen, X., and Dutton, P.L., 1999. Natural engineering principles of electron tunnelling in biological oxidation-reduction, Nature, 402:47–52.

36. Engel, G. S., Calhoun, T. R., Read, E. L., Ahn, T. K., Mancal, T., Cheng, Y. C., Blankeship, R.E., Fleming, G. R., 2007. Evidence for wavelike energy transfer though quantum coherence in photosynthetic systems, Nature, 446: 782–786.

37. Lee, H., Cheng, Y. C., Fleming, G. R., 2007. Coherence dynamics in photosynthesis: protein protection of excitonic coherence, Science, 316:1462–1465.

38. Collini, E., Wong, C. Y., Wilk, K. E., Curmi, P. M., Brumer, P., Scholes, G. D., 2010. Coherently wired light-harvesting in photosynthetic marine algae at ambient temperature, Naure 463:644–647.

39. Ritz, T., Thalau, P., Phillips, J. B., Wiltschko, R., and Wiltschko, W., 2004. Resonance effects indicate a radical-pair mechanism for avian magnetic compass, Nature, 429:177–180.

40. Turin, L., 1996. A spectroscopic mechanism for primary olfactory reception, Chemical Senses, 21, 773–791.

41. McFadden, J., and Al-Khalili, J., 1999. A quantum mechanical model of adaptive mutation, Biosystems, 50, 203–211.

42. Hameroff, S., and Penrose, R., 1996. Orchestrated reduction of quantum coherence in brain microtubules: a model for consciousness, Math. Comp. Simul., 40:453–480.

43. Aharonov, Y., Davidovich, L., Zagury, N., 1993, Quantum random walks, Phys. Rev. A, 48, 1687–1690.

44. Di Molfetta, G., Soares-Pinto, D. O., Duarte Queiros, S. M., 2018, Elephant quantum walk, Phys. Rev. A. 97, 062112.

45. Kurian, P., Dunston, G., and Lindesay, J. 2016. How quantum entanglement in DNA synchronizes double-strand breakage by type II restriction endonucleases, J. Theor. Biology, 391:102–112.

46. Bell, G. I., Jurka, J., 1997, The Length Distribution of Perfect Dimer Repetitive DNA is Consistent with its evolution by an Unbiased Single-Step Mutation Process, J. Mol. Evol. 44, 414–451.

47. Dokholyan, N V., Buldyrev, S. V., Havlin S., Stanley H.E. 2000, Distributions of dimeric tandemrepeats in Non-coding and Coding DNA sequences, J. Theor. Biol. 202, 273–282.

48. Castellanos, M., Mothi, N., Muñoz, V., 2020, Eukaryotic transcription factors can track and control their target genes using DNA antennas, Nature Communications, 11, 540.

49. Jortner, J., Bixon, M., Langenbacher, T., Michel-Beyerle, M. E., 1998, Charge transfer and transport in DNA, Proc. Natl. Acad. Sci, USA, 95, 12759–12765.

50. Camilleri, E., Rohde, P.P., Twamley, J. 2014, Quantum walks with tuneable self-avoidance in one dimension, Sci. Reports, 4, 4791.

51. Bahi, J.M., Guyeux, C., Mazouzi, K., Philippe, L., 2013, Computational investigations of folded self-avoiding walks related to protein folding, Computational Biology and Chemistry, 47, 246–256.

52. Friedman, H., Kessler, D.A., Barkai, E., 2016, Quantum renewal equation for the fisrt detection time of a quantum walk, J. of Physics A: Mathematical and Theoretical, 50.

53. Arrighi, P., Di Molfetta, G., Marquez-Martin, I., Perez, A., 2019, From curved spacetime to spacetime-dependent local unitaries over the honeycomb and triangular Quantum Walks, Sci. Rep. 9 10904

54. Arnault, P., Pepper, B., Perez, A., 2020, Quantum walk in weak electric fields and Bloch oscillations, arxiv:2001.05346.

55. Sapienza P.J., dela Torre, C.A., McCoy IV W. H., Jana, S.V., Jen-Jacobson, L., 2005, Thermodynamics and Kinetic Basis for the relaxed DNA Sequence specificity of Promiscous Mutant EcoRI Endonucleases, J. Mol. Biol, 20, 1–18.

56. Sapienza, P.J., Niu, T., Kurpiewski, M. R., Grigorescu, A., Jen-Jacobson, L., 2014, Thermodynamics and Structural Basis for Relaxation of Specificity in Protein-DNA Recognition, 426,84–104.

57. Stormo, G. D. 2013, Modeling the Specificity of Protein-DNA Interactions, Quant. Biol. 1, 115–130.

58. Etheve, L., Martin, J., Lavery, R. 2017, Decomposing protein-DNA binding and recognition using simplified protein models, Nucleic Acids Research, 45, 10270–10283.

59. Tan C., Takada, S. 2018, Dynamic and Structural Modeling of the Specificity in Protein-DNA Interactions guided by Binding Assay and Structure Data, J. Chem. Theory Comput. 14, 3877–3889.

60. Albuquerque, E.L., Vasconcelos, M.S., Lyra, M.L., de Moura, F.A.B.F 2005, Nucleotide correlations and electronic transport of DNA sequences, Phys. Rev. E, 71, 021910.

61. Lambropoulos, K., Chatzieleftheriou, M., Morphis, A., Kaklamanis, K., Lopp, R., Theodorakou, M., Tassi, M., Simserides, C., 2016, Eletrconicf structure and carrier transfer in B-DNA monomer polymers and dimer polymers, Phys. Rev. E, 94, 062403.

62. Lambropoulos, K., Simserides, C., 2019, Tight-Binding modelling of Nucleic Acid Sequences, Symmetry, 11, 968.

63. Yamada, H., Iguchi, K., 2010, Some Effective Tight-Binding Models for Electrons in DNA Conduction: A Review, Adv. In Condensed Matter Physics, 380710, 1–28.

64. Gans, P.J., 1965, Self-Avoiding Random Walks. I. Simple Properties of Intermediate Length Walks, 42, 4159.

65. van Kampen, N.G., 1998, Remarks on Non-Markov Processes, Brazilian Journal of Physics, 28, 90–96.

66. Vacchini, B., Smirne, A., Laine, E-M., Pillo, J., Breuer, H-P., 2011, Markovianity and Non-Markovianity in quantum and classical systems, New J. Phys. 13, 093004.

67. Roberts, R. J., Vincze, T., Posfai, J., Macelis, D., 2007, REBASE-enzymes and genes for DNA restriction and modification, Nucleic Acids Research 35, D269–D270.

68. Zahran, M., Daidone, I., Smith, J. C., Imhof, P., 2010, Mechanism of DNA Recognition by the Restriction Enzyme EcoRV, J. Mol. Biol. 401, 415–432.

69. Fujitsuka, M., Majima, T., 2017, Charge transfer dynamics in DNA revealed by time-resolved spectroscopy, Chem. Sci., 8, 1752.

70. Wagenknecht, H-A., 2003, Reductive Electron Transfer and Transport of Excess Electrons in DNA, Angew. Chem. Int. Ed., 42, 2454–2460.

71. Garbaczewski, P., Stepanovich, V., 2013, Levy flights and nonlocal quantum dynamics, J. of Mathematical Physics, 54, 072103.

72. Venegas-Andraca, S.E., Ball, J.L., Burnett, K., Bose, S., 2005, Quantum walks with entangled coins, New Journal of Physics, 7, 221.

73. Li, D., McGettrick, M., Zhang, W-W., Zhang, K-J., 2015, One-dimensional lazy quantum walks and occupacy rate, Chinese Phys. B, 24, 050305.

74. Petz D., 2001, Entropy, von Neumann and the von Neumann Entropy. In: Rédei M., Stöltzner M. (eds) John von Neumann and the Foundations of Quantum Physics. Vienna Circle Institute Yearbook, 2000, (Institut ‘Wiener Kreis’ Society for the Advancement of the Scientific World Conception), vol 8. Springer, Dordrecht.

75. Maloyer, O., Kendon, V., 2007, Decoherence versus entanglement in coined quantum walks, New Journal of Physics, 9, 87

76. Mondal, K., Chaudhury, S., 2020, A theoretical study of the role of bulk crowders on target search dynamics of DNA binding proteins, J. Stat. Mech.: Theory and Exp. 093204.

77. Shvets, A.A., Kochugaeva, M.P., Kolomeisky, A.B., 2018, Mechanisms of Proteins Search for Targets on DNA: Theoretical Insights, Molecules, 23, 2106.

78. Mohseni, M., Rebentrost, P., Lloyd S., Aspuru-Guzik, A., 2008, Environment-assisted quantum walks in photosynthetic energy transfer, J. Chem. Phys. 129, 174106.

79. Novo, L., Chakraborty, S., Mohseni M., Omar, Y., 2018, Environment-assisted analog quantum search, Phys. Rev. A, 98, 022316.

80. Berg, O. G., Winter, R.B., von Hippel, P.H., 1981, Diffusion-driven mechanisms of protein translocation on nucleic acids. 1. Models and theory, Biochemistry, 20, 6929–6948.

81. Bhattacherjee, A., Levy, Y., 2014, Search by proteins for their DNA target site: 1. The effect of DNA conformation on protein sliding, Nucleic Acids Research, 42, 12404–12414.

82. Mahmutovic, A., Berg, O. G., and Elf, J. 2015. What matters for *lac* repressor search *in vivo*-sliding, hopping, intersegment transfer, crowding on DNA or recognition?, Nucleic Acids Res. 43:3454–3464.

83. Alberti, A., Alt, W., Werner, R., Meschede, D., 2014, Decoherence models for discrete-time quantum walks and their application to neutral atom experiments, New J. of Physics, 16, 123052.

84. Fahri, E., Goldstone, J., Gutmann, S, 2008, A quantum algorithm for the Hamiltonian NAND tree, Theory Comput. 4, 169–190.

85. Wang, Y., Cui, Z-W., et al, 2020, Quantum slide and NAND Tree on a Photonic Chip, arXiv:2001.08755v2.

86. Agliari, E., Blumen, A., Mülken, O., 2010, Quantum-walk approach to searching on fractal structures, Phys. Rev. W, 82, 012305.

87. Kempe, J., 2003, Quantum random walks: an introductory overview, Contemp. Phys. 44, 307–327.

88. Childs, A.M., 2010, On the relationship between continuous- and discrete-time quantum walk, Communications in Mathematical Physics, 294, 581–603.

89. Childs, A. M., Fahri, E., Gutmann, S., 2002, An example of the difference between quantum and classical random walks, Quantum Information Processing, 1, 35–42

90. Gradshteyn, I.S., Ryzhik, I.M., 2007 Table of Integrals, Series and Products, Seventh English version, Academic Press.

91. Mülken, O., Blumen, A., Amthor, T., Gise, C., Reetz-Lamour, M., Weidemüller, M., 2007, Survival Probabilities in Coherent Transfer with Trapping, Phys. Rev. Lett. 99, 090601.

92. Chupeau, M., Bénichou, O., Voituriez, R., 2015, Cover times of random searches, Nature Physics, 11.

93. Childs, A.M., Goldstone, J., 2004, Spatial search by quantum walk, Phys. Rev. A, 70, 022314.

